# Deep mutational scanning of EccD_3_ reveals the molecular basis of its essentiality in the mycobacterium ESX secretion system

**DOI:** 10.1101/2024.08.23.609456

**Authors:** Donovan D. Trinidad, Christian B. Macdonald, Oren S. Rosenberg, James S. Fraser, Willow Coyote-Maestas

**Affiliations:** Department of Medicine, University of California, San Francisco; Department of Bioengineering and Therapeutic Sciences, University of California, San Francisco, San Francisco, CA 94158, USA; Quantitative Biosciences Institute, University of California, San Francisco, USA; Chan Zuckerberg Biohub, San Francisco, CA, USA

## Abstract

Tuberculosis remains the deadliest infectious disease in the world and requires novel therapeutic targets. The ESX-3 secretion system, which is essential for iron and zinc homeostasis and thus *M. tuberculosis* survival, is a promising target. In this study, we perform a deep mutational scan on the ESX-3 core protein EccD_3_ in the model organism *M. smegmatis*. We systematically investigated the functional roles of 145 residues across the soluble ubiquitin-like domain, the conformationally distinct flexible linker, and selected transmembrane helices of EccD_3_. Our data combined with structural comparisons to ESX-5 complexes support a model where EccD_3_ stabilizes the complex, with the hinge motif within the linker being particularly sensitive to disruption. Our study is the first deep mutational scan in mycobacteria, which could help guide drug development toward novel treatment of tuberculosis. This study underscores the importance of context-specific mutational analyses for discovering essential protein interactions within mycobacterial systems.

## Introduction

*Mycobacterium tuberculosis*, the causative agent of tuberculosis, poses a significant global health threat, claiming the lives of over a million people annually^1^. The innate immune response to intracellular pathogens is complex, including strategies such as starvation by phagocytosis and sequestration of essential trace metals in a process called nutritional immunity. *M. tuberculosis* is able to survive and establish replicative niches in the host through the release of numerous proteins through specialized secretion systems, such as the 6 kDa early secretory antigenic target (ESAT-6) protein family secretion (ESX) systems. There are five paralogous ESX secretion systems (ESX-1 to -5) that play unique roles in virulence and nutrient acquisition^2–8^. ESX-3 in particular is an attractive therapeutic target because it is involved in iron and zinc homeostasis, making it essential for *M. tuberculosis* survival *in vitro* and *in vivo*^8–10^. Development of such therapies requires an understanding of the ESX-3 complex and how it functions.

Structures of the ESX systems provide key information on which proteins make up the core complex and how the secretion system assembles. Two independently determined structures of the ESX-3 complex, purified from the model organism *M. smegmatis,* revealed a dimeric complex that extends into both the periplasm and cytoplasm^11,12^. Each protomer complex consists of one copy of EccB_3_, EccC_3_, and EccE_3_, and a dimer of two asymmetrical copies of EccD_3_, where there are notable interfaces between the soluble domains of EccC_3_, EccD_3_, and EccE_3_ (Fig 1A, B). Intriguingly, there are also two high-resolution hexameric structures of the homologous ESX-5 complex that retain the same core complex architecture, but are fully assembled as a trimer of subcomplex dimers^13,14^. Despite this structural information, the molecular basis of how ESX-3 recognizes and secretes substrates remains unclear.

**Figure 1.**
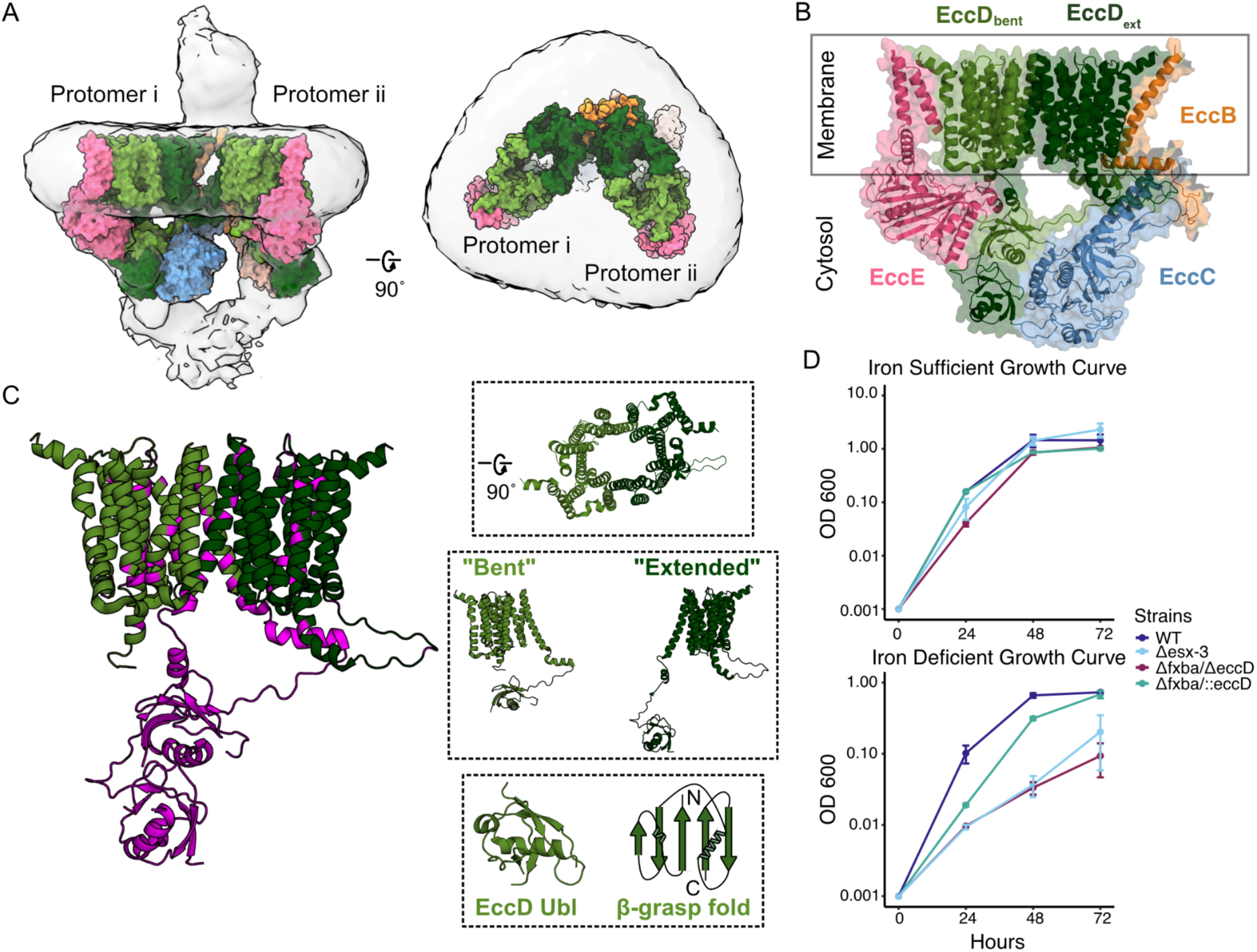
The Ubl domain, conformationally distinct linker, and vestibule make the EccD protein an attractive target. A) The ESX-3 dimer complex, where the model PDB: 6UMM is fit in the EM map EMDB: 20820. Surface representation of the ESX-3 model showing two protomers, with one copy per protomer of EccB3 (orange), EccC3 (blue), EccE3 (pink), EccD3-bent (light green), and EccD3-extended (dark green). Left: Front view of the ESX-3 dimer. Right: Top down view of the ESX-3 dimer, highlighting the open vestibule the EccD dimers form. Modeled using ChimeraX. B) Surface and cartoon model of one ESX-3 protomer. C) View of the EccD3 dimer, where residues targeted in this study are colored purple. Top inset: View of the EccD vestibule. Middle inset: The two EccD conformational states, “bent” and “extended”. Bottom inset: View of the EccD ubiquitin-like domain, and a cartoon representing the β-grasp fold shared with ubiquitin. D) Growth curves showing the previously observed low-iron growth phenotype. The growth of the wild-type, Δ*fxbA*/Δ*eccD3*, Δ*fxbA/::eccD3*, and *Δesx-3* strains was monitored by determining their optical densities at 600 nm. The experiments were performed 3 times and error bars represent the standard deviations across replicates.

We were particularly interested in studying EccD_3_ due to its position within the ESX complex, as EccD_3_ interacts with each of the other core proteins both in the membrane and the cytosol, creating many large interfaces that could theoretically be targeted by a small molecule and disrupted to abolish function (Fig 1C). The EccD_3_ dimer interacts within the membrane to form a large vestibule filled with lipids. There is a flexible linker that is positioned in two different conformational states across the EccD_3_ dimer, which connects the first transmembrane helix to the soluble ubiquitin-like (Ubl) domain. Ubiquitin and Ubl proteins and domains have an evolutionarily conserved β-grasp fold where a beta sheet composed of five anti-parallel beta strands are arranged around (or “grasp”) a single alpha helix^15,16^. Ubl proteins and domains serve as scaffolds across a variety of biological contexts, and often play different functional roles than ubiquitin, especially in bacteria that do not contain a eukaryotic-like ubiquitin-protease system^17–19^. EccD_3_ is also essential to ESX-3 complex function, and organism survival^20^ (Fig 1D).

The expansion of the mycobacteria genetic toolkit alongside advances in molecular biology allows us to utilize novel approaches to identify druggable targets. Tools such as TnSeq and CRISPR enable high-throughput functional genomic screens to identify and annotate genes with similar phenotypes through gene knockout or knockdown^21–25^. These screens reveal essentiality of protein systems, including ESX-3, however they fall short in elucidating underlying mechanisms and specific interactions. Deep mutational scanning (DMS) is a powerful technique that addresses this gap and allows us to explore intricate amino acid interaction networks beyond structural insights. Mutational scanning facilitates a comprehensive analysis of residue essentiality by creating libraries where wild-type amino acids are systematically replaced with each of the other 19 amino acids, and then screened via selection to reveal interactions crucial for structure or function^26^. After the identification of a potential drug target, this approach can guide the development of therapeutics that target specific interactions.

We performed DMS on EccD_3_ to determine how residues contribute to function and generate hypotheses on its overall role within the ESX-3 complex. We used *M. smegmatis*, a model organism for *M. tuberculosis*, and leveraged *M. smegmatis’* dependence on ESX-3 for iron acquisition to design a high-throughput growth-based selection assay to screen EccD_3_ variants. We made comparisons to a prior DMS on ubiquitin to increase signal-to-noise in our interpretation of the Ubl domain mutagenesis data. Together this data provides insight into how EccD_3_ supports the ESX-3 complex, and combined with our structure, allow us to identify residues important for folding & stability, residues important for mediating protein-protein interactions seen in the structure, and residues that may participate in switching between oligomeric states.

## Results

### A fxbA/eccD_3_ double knockout strain provides an iron sensitive conditional selection for EccD_3_ function

To enable functional selections of EccD_3_ mutant variants, we adapted a growth assay based on the established low-iron growth phenotype in *M. smegmatis*^10,20^. This selection relies on the requirement of a functioning ESX-3 for iron acquisition mediated by the mycobactin siderophore. Knockout of the entire ESX-3 operon leads to inhibited *M. smegmatis* growth in a low-iron environment. When individual components of the ESX-3 system are deleted, growth is only impaired if the additional siderophore exochelin formyltransferase fxbA is also knocked out^20^. Therefore, our selection used a strain lacking the endogenous *eccD_3_* and *fxbA* genes. To validate the selection, we measured growth in chelated Sauton’s media (iron-deficient) and in Sauton’s media with iron spiked in (iron-sufficient) for three days, comparing the double knockout of fxbA and eccD_3_, (Δ*fxbA/*Δ*eccD_3_*) with complement of wild-type (WT) eccD_3_ in the fxbA knockout background (Δ*fxbA/::eccD_3_*). As expected, the complemented strain behaves like WT *M. smegmatis* whereas the uncomplemented strain behaves similarly to a knockout of the entire ESX-3 operon (*Δesx-3*) (Fig 1D). While all strains grew equally as well in the iron-sufficient media, a stark growth defect was noted in the Δesx-3 and Δ*fxbA/*Δ*eccD_3_* strains in the iron-chelated media, whereas complementation with full length EccD_3_ rescued growth, validating the premise of the selection.

### A mutagenesis library for EccD_3_ functional selections

We reasoned that if our mutational scan revealed that most residues in which mutations are deleterious play scaffolding roles, this would imply a structural role supporting the other core ESX-3 components. In contrast, if we found an enrichment of residues that extend into the vestibule, this would imply EccD_3_ plays a role more directly related to ESX-3 activity, perhaps acting as a pore to secrete substrates. To test these hypotheses, we generated a saturation mutagenesis library for selected positions to understand how EccD_3_ contributes to ESX-3 function. Our library comprised the complete cytosolic ubiquitin-like (Ubl) domain and the flexible linker that connects the Ubl domain to the transmembrane domain (Fig 1C). We only selected residues in the transmembrane domain whose side chains extend into the vestibule or are conserved across all mycobacterial ESX systems, as we were limited by synthetic constraints. For each of these residues, our library of variant oligonucleotides contained all codon optimized amino acid mutations, a stop codon, and (where possible) a synonymous WT codon. We designed a library of 3965 variants (165 residues x 21), however we received a pool of 3045 variants from Twist Bioscience as 20 sites failed in DNA synthesis. The reduced positional coverage in these regions is likely due to high GC content limiting synthesis and quality control. We introduced the 3045 variant library into the pMV306 plasmid which contains the mycobacteriophage L5 attP attachment site, which allows for stable integration into the mycobacterial genome at the chromosomal attB insertion site (Fig 2A)^27,28^.

**Figure 2.**
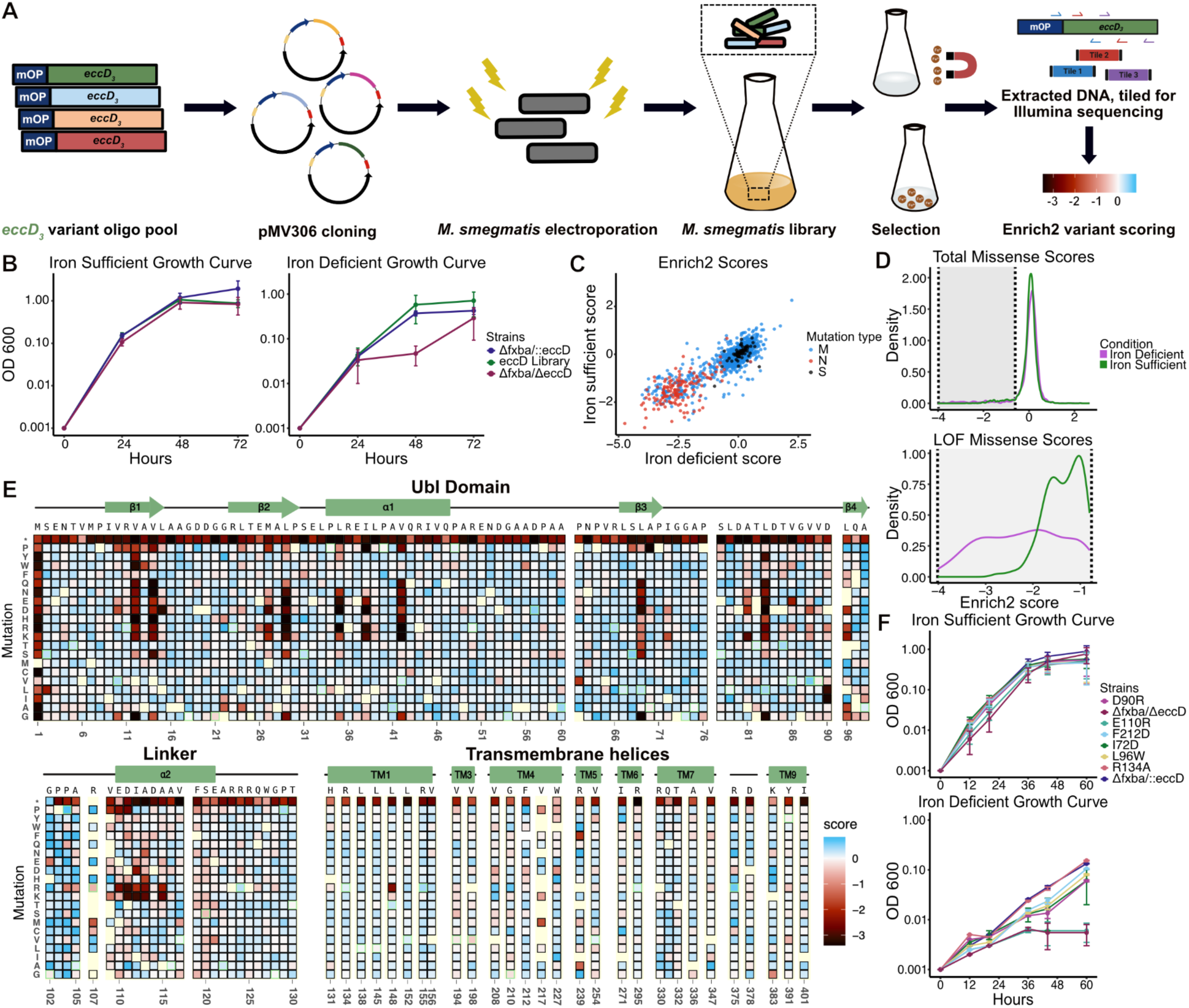
Deep mutational scan of the EccD protein reveals residues critical for function. A) Schematic of the EccD variant library generation and growth assay. A mutant variant library was designed to include the mycobacterial optimized promoter (mOP) and ordered as an oligo pool. The library was cloned into the integrative mycobacterial plasmid pMV306 and electroporated into *M. smegmatis*. Growth of the *M. smegmatis* library in low- or high-iron media was monitored and samples were collected every 24 hours. PCR was performed on extracted DNA to generate overlapping amplicons for deep sequencing. The sequencing data was analyzed and assigned a functional score using Enrich2. B) Growth of the positive control Δ*fxbA/*Δ*eccD3* strain, negative control Δ*fxbA/::eccD3* strain, and the EccD3 *M. smegmatis* library in the chelated iron-deficient Sauton’s or the iron-sufficient Sauton’s medium. The experiments were performed 3 times and error bars represent the standard deviations across replicates. C) Scatter plot of the functional scores of each variant when grown in iron-sufficient versus iron-deficient conditions. Missense mutations are colored blue, nonsense mutations are red, and synonymous mutations are black. D) Density plots representing the distribution of Enrich2 scores for missense variants in both conditions. Top: Total distribution of missense scores in both conditions. Bottom: Distribution of loss-of-function missense scores in both conditions. E) Heatmap of the EccD3 functional scores from the iron-deficient growth condition. The WT sequence, domain organization, and cartoon secondary structure representation of EccD3 are shown above each section of the heatmap. The variant identity is indicated on the y-axis, and the residue position is indicated on the x-axis. WT-synonymous substitutions are outlined in green and missing data are light yellow. F) Growth of selected point mutants in the chelated iron-deficient Sauton’s or the iron-sufficient Sauton’s medium was monitored by determining their optical densities at 600 nm. The experiments were performed 2 times and error bars represent the standard deviations across replicates.

In order to screen our EccD_3_ variants in the native cellular environment, we next needed to electroplate our plasmid library into the Δ*fxbA/*Δ*eccD_3_* strain. Our initial attempts at electroporation resulted in substantially fewer colonies than required for a saturated library. We optimized the procedure by modifying the volume of electrocompetent cells, the concentration and volume of input DNA, and recovery intervals. Our electroporation efficiency increased from 7.6x10^4^ to 6x10^5^ transformants/µg after optimization (Table 1). Recombinants were selected on kanamycin-containing plates and showed stable integration by colony PCR and Sanger sequencing. Deep sequencing prior to selection revealed that our final library in *M. smegmatis* contained 2972 variants (97.6% the received library).

### DMS experiments reflect EccD_3_ functional requirements in the presence and absence of iron

To identify residues essential to ESX-3 function, we grew *M. smegmatis* cells bearing our library in the presence of iron (iron-sufficient media), and the absence of iron (iron-deficient media). We collected samples across three replicates at 24 hour timepoints to measure OD600 and to extract DNA for sequencing (Fig. 2B). We used Enrich2 to calculate variant fitness scores that represent changes in mutation frequency relative to WT throughout the experiment, for both conditions^29^. Positive scores reflect an increase in frequency throughout the experiment and increased growth relative to WT, while negative scores reflect a decrease in frequency and a deleterious fitness effect. Based on the growth curves in Fig. 1, we expected the variant library to have scores similar to WT when grown in the iron-sufficient media, and for deleterious effects to emerge in the iron-deficient media.

We observed that most mutations have similar scores in both conditions (Fig 2C). Synonymous and the majority of missense mutations (94.19% in iron-sufficient media; 93.36% in iron-deficient media) cluster together and display near WT fitness, reflecting tolerance to mutations. Nonsense mutations were highly shifted away from the main population of WT and missense mutations in both conditions, and have strong loss-of-function (LOF) effects. We observe that iron starvation sensitizes deleterious effects seen in premature stop codons and some missense mutations further, leading to fitness scores being more pronounced in the iron-deficient media (Fig 2D). Of the 270 LOF mutations seen in the iron-deficient condition, 37 (13.7%) were tolerant in the iron-sufficient condition, and 39 (14.44%) had strong LOF effects but weak LOF effects in the iron-sufficient condition. Conversely, growth in the more permissive iron-sufficient medium reflected a more mutationally tolerant environment. Gain-of-function (GOF) mutations are rare in both conditions (9/2987 (0.3%) in iron-deficient media, 41/2987 (1.37%) in iron-sufficient). There is also a small set of residues (V208, G210, F212, V217, W227) where missense mutations are tolerated in the iron-deficient media but deleterious in the iron-sufficient media, suggesting these substitutions are more impactful when iron is readily available in the environment.

Based on our growth curve analysis, we expected the mutant variant scores to reflect a more tolerant environment when grown in the iron-sufficient media. That we did not see emergent phenotypes in the iron-deficient condition implies EccD_3_ is important for growth even when there is an increased amount of iron in the environment. This speaks to the sensitivity of next-generation sequencing compared to the strong differences observed between conditions in phenotypic growth curves. However, the removal of iron sensitizes the environment and leads to a broader range of effects when the variant library is studied in the iron-deficient media (Fig 2D). As we observed that mutations were largely important under both conditions, we proceeded with analysis in the iron-deficient condition (Fig 2E).

### Individual growth curves recapitulate library growth rates

To validate the results of the DMS pooled growth rates, we screened a small library of point mutants (Fig 2F). We chose 7 residues with a range of functional effects from each domain, where 2 mutants reflected tolerant substitutions, 4 mutants had weak LOF effects, and 1 mutant had a strong LOF effect. We screened: I72D, D90R, L96W within the Ubl domain, where I72 and D90 are residues at protein-protein interfaces, and L96 is a buried, hydrophobic residue within the domain core; E110R and R134A within the flexible linker, where E110 maintains salt-bridge interactions at interfaces, and R134 is at the end of the linker; and F212D, where F212 is in the periplasmic loop connecting transmembrane helices 3 and 4. R134A was entirely tolerant to mutation, as seen in our heatmaps. By comparison, I72D, D90R, L96W, and F212D had weak LOF effects, and E110R had a strong LOF effect. These effects are reflected within the growth curve, where R125A grows at faster rates than the other mutants and is closest to the growth rate of the WT complemented strain. I72D, D90R, and L96W grow slightly slower than the WT. E110R has the most dramatic phenotype, and grows as poorly as the double knockout strain. Thus, we see that our deep mutational scan data accurately represent *M. smegmatis* growth.

### A stably folded ubiquitin-like domain is required for ESX-3 function

Next, we examined the mutational tolerance of the N-terminal 99 amino acid (aa) ubiquitin-like (Ubl) domain of EccD_3_, which form interfaces between the soluble domains of each EccD_3_ monomer, as well as EccB_3_, EccC_3_, and EccE_3_ in the ESX-3 dimer structure (PDB: 6UMM). Analyzing the mutational data within the EccD_3_ Ubl domain reveals most missense mutations are tolerated (92.76%). There were only strong LOF effects for the hydrophobic core residues (V12, V14, M27, L29, L35, I38, V42, L69, L83, and L96), which are tolerant only to hydrophobic substitutions (Fig 3A, B). A81 is intolerant to swaps to the charged residues D, E or K, and is in a loop connecting beta sheets 3 and 4, near the buried core residues L35, I38, and L83 which may explain this sensitivity. These patterns were observed in both growth conditions.

**Figure 3.**
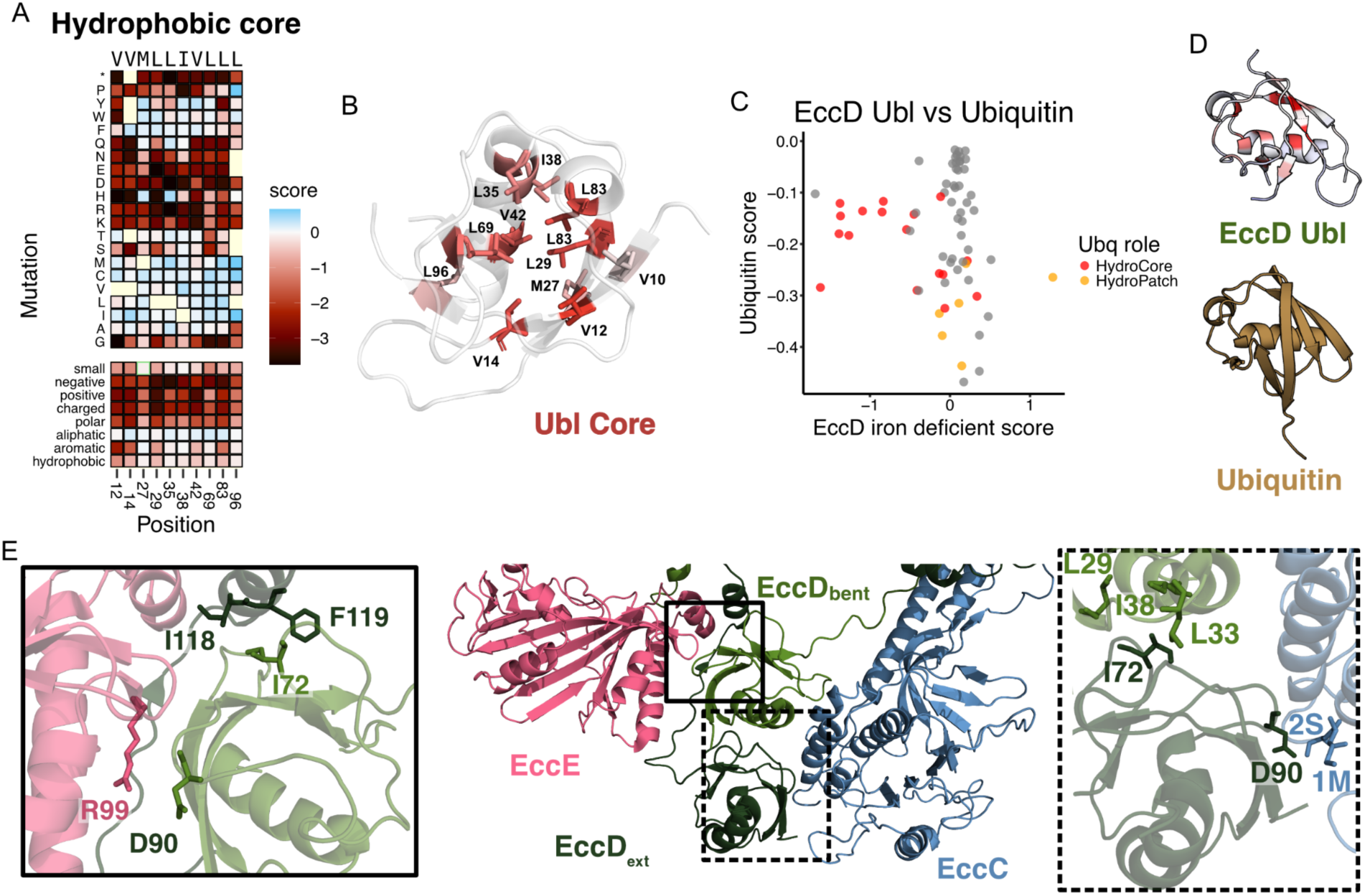
The EccD3 ubiquitin-like domain is sensitive to mutations in the hydrophobic core. A) Subset heatmap of the Ubl domain hydrophobic core from the iron-deficient selection. B) The mean missense mutation functional scores mapped onto the Ubl domain hydrophobic core residues, from low (red) to white (tolerant) scores. All other residues are colored white. C) Scatter plot of mutational effects seen at a given position in a DMS of ubiquitin versus the DMS of Ubl domain. Residues that make up the ubiquitin hydrophobic core are colored red, residues that make up the ubiquitin hydrophobic patch are colored orange, and all other residues are grey. D) The EccD Ubl domain and ubiquitin (PDB: 1UBQ), where the Ubl domain is colored by average missense mutation scores from the iron-deficient growth condition. E) The EccD Ubl domain interfaces with each other and the core proteins EccE and EccC. Left inset: residues highlighted at the EccD3-bent Ubl domain interface with EccE. Right inset: residues highlighted at the EccD3-ext Ubl domain interface with EccC. EccE colored pink, EccC colored blue, EccD3-bent colored light green, and EccD3-ext colored dark green.

Comparisons to prior DMS screens on ubiquitin can help distinguish which residues are required to maintain stability of the β-grasp fold from residues that are unique to the function of EccD_3_ in the ESX-3 complex^30–32^. To contextualize the results from our screen, referenced prior screens of ubiquitin in yeast where growth was linked to ubiquitin-mediated protein degradation (Fig 3C, D). In ubiquitin, the strongest LOF effects were in residues that performed well-established roles in E1 activation and poly-ubiquitin linkage formation, and residues within the surface hydrophobic patch^31^. The hydrophobic core residues all are sensitive to swaps to charged residues, but those that are near functionally essential residues are far more sensitive to polar substitutions with the strongest LOF effects clustered near the β-sheet surface and C-terminus^31^. In contrast to ubiquitin, all hydrophobic core residues in the EccD_3_ Ubl domain are equally intolerant to charged residue swaps. Unsurprisingly, residues important for ubiquitin’s specific degradation interactions are not sensitive to substitutions in the EccD_3_ Ubl domain. There is no simple discernable preference within the Ubl domain to any side that maintains protein-protein interactions, implying that the scores are dominated by stability effects.

The EccD_3-bent_ Ubl domain forms interfaces with the EccD_3-ext_ Ubl domain as well as the EccC_3_ and EccE_3_ soluble domains (Fig 3E). The EccD_3-ext_ Ubl domain forms interfaces with the EccD_3-bent_ Ubl domain and EccC_3_. Most EccC_3_ and EccE_3_ interface residues tolerate substitutions apart from I72 and D90. The I72 site is analogous to the primary protein-protein interaction site in ubiquitin: the ubiquitin hydrophobic patch, which is made up of residues I8, I44, and V70, all of which are highly sensitive to mutation. While the analogous residues in EccD_3_ are hydrophobic, this patch is tolerant to mutation in EccD_3_, in contrast to ubiquitin. The exception is I72, which has LOF effects when swapped to aspartic acid or lysine. In EccD_3-ext_, I72 is at the hydrophobic interface between the two Ubl domains, extending toward the EccD_3-bent_ hydrophobic core. In EccD_3-bent_, I72 is in a pocket of hydrophobic residues between EccE_3_ and the EccD_3-ext_ linker. A swap from the hydrophobic isoleucine to a charged residue likely disrupts the EccD_3-bent_:EccD_3-ext_ interface, and displaces the EccD_3-ext_ linker. D90 is the other interface residue that is sensitive to mutation, and is on a loop between the 4th and 5th beta strands of the Ubl domain. This position is buried at the interfaces between EccE and EccD_3-bent_, and EccC and EccD_3-ext_. While a charge-swap substitution to an arginine has a weak LOF effect, substitutions to the hydrophobic isoleucine or leucine have strong LOF effects. This implies the salt-bridge interaction between EccD_3-bent_ D90 and EccE_3_ R99 is not necessary to ESX-3 function, as a charge swap does not affect growth. In contrast, a swap to a branched hydrophobic is deleterious. Together, this data suggest that stability of the EccD_3_ Ubl domain is important for ESX-3 function, while interactions between key residues at the interfaces with neighboring proteins are essential under the conditions we assayed.

### The EccD_3_ cytosolic linkers maintain salt bridge interactions between other ESX-3 core components

A cytosolic linker (residues 100-130) connects the soluble Ubl domain to the first transmembrane helix and establishes the two placements (bent and extended) of the Ubl domain in the dimeric structure. The mutational landscape reveals the strongest signals at the start of the linker: G102, P103, P104, and A105 (Fig 4A). These residues are tolerant to mutation and, surprisingly, also display GOF phenotypes in iron-sufficient media. This pattern suggests that conformational requirements for Gly and Pro within the linker are not needed in this growth condition. Despite observing high fitness values upon mutation, this region of the linker is important for allowing the distinct asymmetrical conformations observed in each EccD_3_ monomer. In EccD_3-ext_, the linker extends alongside the EccE_3_ and EccD_3-bent_ interface, and folds into a short α-helix before beginning the first transmembrane helix (Fig 4C-F). In contrast, in EccD_3-bent_, the linker runs alongside the EccC_3_ stalk domain and “bends” before folding into an α-helix that contacts EccB_3_ and EccC_3_ in a hinge region between each ESX protomer (Fig 4C, G-I). This bend reorients the linker in the opposite direction so that it can fold adjacent to the first transmembrane helix. Given these distinct residue environments, we investigated whether the pattern of mutations could distinguish a stronger functional requirement for either conformation in other regions of the linker.

**Figure 4.**
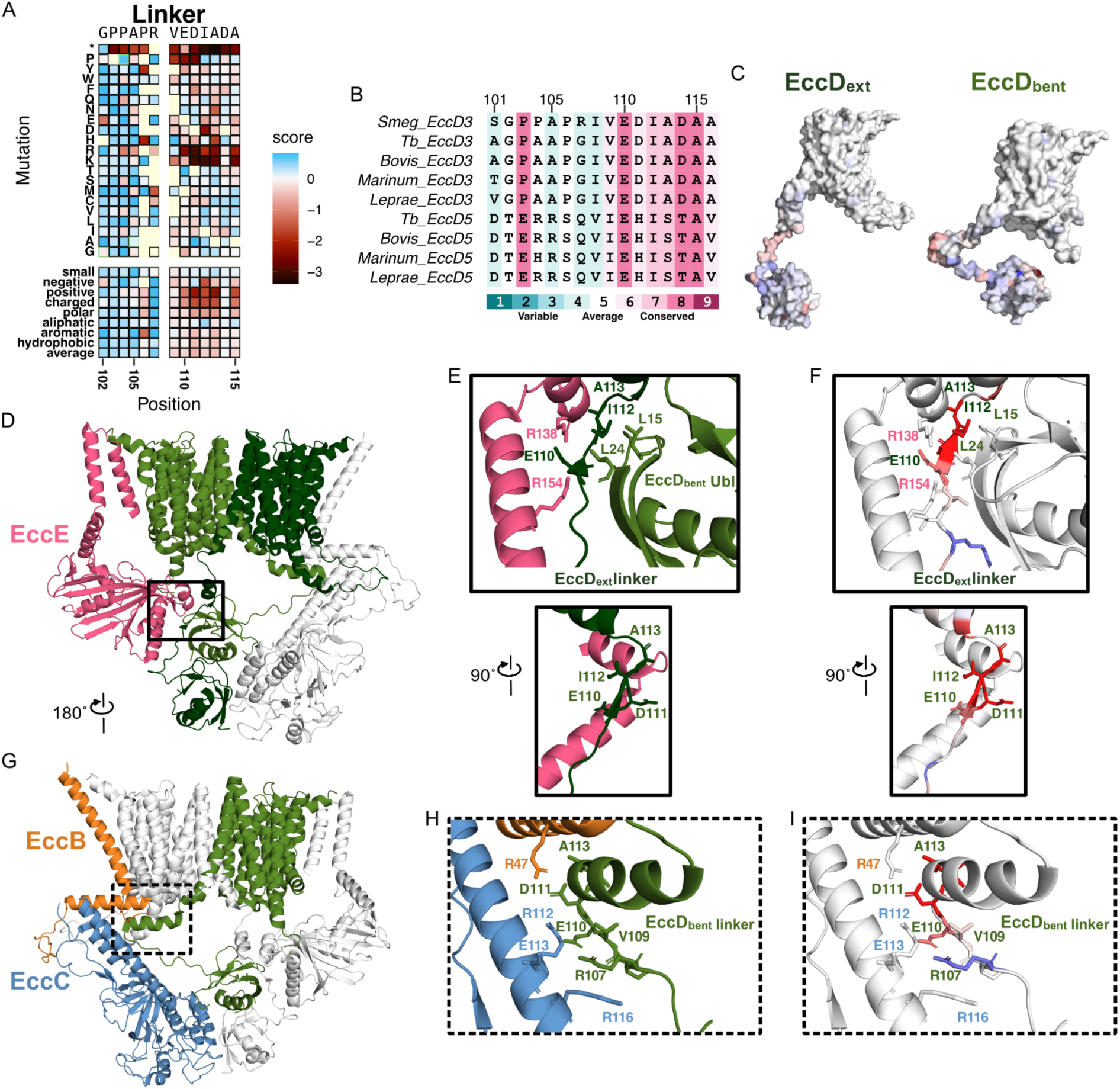
The EccD3 linker is sensitive to positive-charge mutations. A) Heatmap section of the linker domain from the iron-deficient selection. B) Multiple sequence alignment of *M. smegmatis* EccD_3_ and homologs, colored by conservation scores on a gradient where blue represents low conservation, white represents average conservation, and red represents high conservation. Performed using Consurf^33^. C) The mean missense mutation functional scores mapped onto the EccD3-ext and EccD3-bent monomers. D) Front view of the ESX-3 protomer highlighting the EccD3-ext linker in the solid box. EccE colored pink, EccD3-bent colored light green, and EccD3-ext colored dark green, and EccC and EccB colored white. E) The EccD3-ext linker interactions with EccE. Top: Front view of the interaction. Bottom: Side view of the interaction to highlight D111. F) The same views as E, with the average positive-mutation functional scores mapped onto the EccD3-ext linker residues. EccE is colored white. G) Back view of the ESX-3 protomer highlighting the EccD3-ext linker interaction with EccB and EccC in the dotted box. EccB colored orange, EccC colored blue, EccD3-bent colored light green, and EccD3-ext and EccB colored white. H) The EccD3-bent linker interactions with EccB and EccC. I) The same view as H, with the average positive-mutation functional scores mapped onto the EccD3-bent linker.

Examining the heat map from the point of view that one conformational state may be more functionally important helps interpret the patterns for highly conserved residues E110, D111, I112, A113, D114, and A115, which have strong LOF when mutated to positively charged (arginine or lysine) residues (Fig 4B). Additionally, E110 and D111 have strong LOF effects when swapped to a proline. In EccD_3-bent_, these residues form a charged network of stabilizing interactions with the EccC stalk domain and an EccB_3_ membrane helix, and is adjacent to the loop between EccD_3-ext_ transmembrane helices 6 and 7 (Fig 4F). This forms a hinge region, where the two ESX complexes interact to form the dimer complex. The strong LOF effect from charge swapping the E110 and D111 residues suggests the salt-bridge interactions between E110 and R112, and D111 and R47 are required for ESX-3 function. This extends to A113, where swapping to an arginine or lysine would disrupt the interaction between D111 and R47. It is surprising that D114 tolerates all substitutions despite extending into the same pocket. On the other side of the EccD_3-bent_ helix, I112 and A115 extend into a hydrophobic pocket created by the loop between EccD_3-ext_ TM 6 and TM 7. In EccD_3-ext_, the linker is positioned at the back of the EccD_3-bent_ Ubl and EccE_3_ soluble domain interface (Fig 4E). Similarly to the EccD_3-bent_ linker, E110 forms a salt bridge with the nearby R138 on EccE. In contrast, D111 does not form any interaction at the PPI, and instead extends into cytosolic space. The residues I112, A113, and A115 are in a hydrophobic patch with the EccD_3-bent_ Ubl residues L15 and L24. Introducing a positive charge would disrupt this interface, and I112R specifically would clash and be electrostatically incompatible with EccE_3_ R138. This data suggests the linker domain must be able to form stabilizing contacts with the other ESX-3 core proteins to maintain the core complex architecture, and that the sensitivity to mutation may reflect the importance of the EccD_3-bent_ in stabilizing the ESX dimer.

### The 3rd and 4th transmembrane helices display sensitivity in the iron-sufficient condition

In the EccD_3_ dimer, each EccD_3_ monomer has 11 transmembrane helices where transmembrane helix 1 in one EccD_3_ interacts with transmembrane helices 9 and 10 in the next EccD_3_ to form a large vestibule that measures approximately ∼20x30 Å in cross-sectional diameter. In the ESX-3 structures, the periplasmic half of the vestibule is lined with eight densities that are consistent with lipid tails or detergent molecules. To study whether these densities were functionally important and identify other residues that may have importance, we chose 40 residues for saturation mutagenesis based on side chain contacts with lipid densities and evolutionary conservation. We successfully measured the effects of some substitutions in 35 and most substitutions (at least 10/20 missense mutations) in 29 out of an intended 40 residues.

We found that 98.73% of transmembrane domain variants are tolerated when grown in iron-deficient media, compared to 96.73% of variants grown in iron-sufficient media. The residues that extend from the transmembrane helices out into the vestibule tolerate substitutions. The exception is residue T148, which is largely tolerant to missense mutation but shows a strong LOF effect when swapped to an arginine in both growth conditions. T148 extends out from transmembrane helix 1 into a hydrophobic pocket between transmembrane helices 1, 2, and 3. A substitution to arginine would not extend into the lipid channel itself, however it may clash with transmembrane helix 3. That we don’t see functional consequences for substitutions implies these lipids are not important for the screened iron acquisition phenotype.

Our heatmaps surprisingly reveal residues where substitutions are deleterious specifically in the iron-sufficient condition (Fig 5A, B). In the iron-deficient condition, 6/551 (1.08%) missense mutations have a weak LOF effect, and 0 have strong effects. In the iron-sufficient condition, 8/551 (1.45%) missense mutations have a weak LOF effect and 6/551 (1.08%) have strong LOF effects. The residues that have these unique effects are V208, G210, F212, V217, W227, and R239. V208, G210, and F212 are in the periplasmic loop connecting transmembrane helices 3 and 4. Swaps to negatively charged residues have weak LOF effects, while in V208 swaps to a positively charged arginine or lysine have GOF effects. V217 is in transmembrane helix 4, and may interact with one of the vestibule lipids. W227 extends into a hydrophobic pocket between transmembrane helices 3 and 4. In the iron-deficient condition, swaps to proline, tryptophan, and aspartic acid have a weak LOF effect, whereas most other swaps are tolerant or have weak GOF effects. In contrast, swaps to proline or tryptophan have the strongest LOF effects in the iron-sufficient condition, followed by swaps to negatively charged residues, while other swaps remained tolerant or GOF. R239 is at the bottom of transmembrane helix 5 and extends into the membrane channel, and most substitutions are deleterious implying it plays an important role in ESX function. In iron chelated media swaps to phenylalanine and alanine are deleterious, and in iron-sufficient media this also includes aspartic acid and glutamic acid. It is not immediately clear what functional role these residues play.

**Figure 5.**
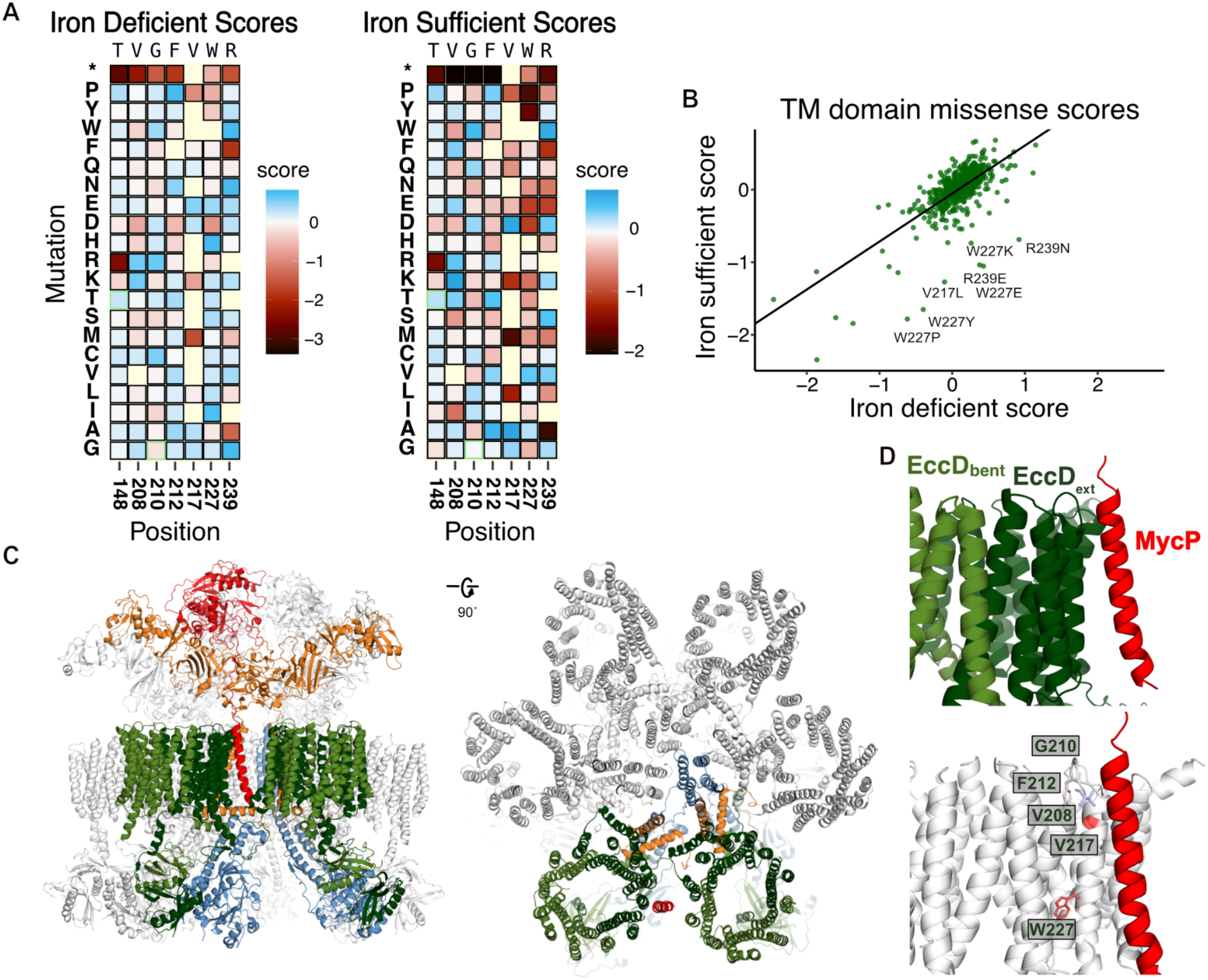
The EccD 3rd and 4th transmembrane helices may play a role in MycP system-specificity. A) Subset heatmap of key residues in TM domain from the iron-deficient and iron-sufficient growth conditions. B) Scatter plot of the missense mutation functional scores from the iron-deficient and iron-sufficient conditions. C) Structure of the ESX-5 complex purified from *M. tuberculosis* (PDB: 7NP7). Left: Front view of the complex. Right: Rotated 90 degrees to reach a top-down view of the complex. One subcomplex is colored the same as ESX-3, the rest are colored white. EccD5-bent colored light green, EccD5-ext colored dark green, EccC5 colored blue. EccB5 colored orange, MycP5 colored red. D) View of the ESX-3 EccD3 dimer with MycP5 modeled in. Top: view with the proteins colored the same as C. Bottom: EccD3-ext is colored white, with the residues with the emergent phenotypes in the iron-sufficient condition colored by DMS score. V208, G210, F212, and V217 are colored by the average positive score, W227 is colored by the average negative score.

When examining the ESX-3 dimeric structure, these residues do not interact with other core proteins, and do not appear to affect complex assembly. However new interactions are revealed when we superimpose the ESX-3 dimer onto the ESX-5 hexamer structures (PDB: 7NP7) (Fig 5C). In the ESX-5 hexamer, three ESX-5 dimer subcomplexes assemble with transmembrane helical rearrangements to allow for this oligomerization^13,14^. The *M. tuberculosis* ESX-5 hexamer was purified with the accessory protein MycP_5_ still bound^13^. Notably, the MycP_5_ transmembrane helix inserts into the membrane alongside the EccD_5-ext_ transmembrane helix 4, and continues into the pocket formed between the EccD_5-ext_ transmembrane helix 11 and EccD_5-bent_ transmembrane helix 1 in the next ESX dimer subcomplex (Fig 5D). In the ESX-3 complex, MycP_3_ may insert near the hydrophobic residues extending from transmembrane helix 4, including V208, G210, and F212 in the periplasmic loop. The MycP transmembrane helix has been hypothesized to be required for ESX complex specificity, targeting MycP to associate with the correct ESX homologue. The effects of mutations near the MycP binding site in ESX-5 homologs hints potentially at these residues playing key roles in maintaining periplasmic interactions, and more broadly a role of EccD_3_ in ESX-3 hexamer complex assembly.

## DISCUSSION

We performed a deep mutational scan of the ESX-3 complex core protein EccD_3_ by growing mutants in iron-deficient and iron-sufficient conditions. We assembled a library to measure the effects of all possible single amino acid substitutions in 145 residues that comprise the ubiquitin-like (Ubl) domain, the flexible linker that connects the Ubl domain to the first transmembrane helix, and a subset of residues across the transmembrane helices. In the Ubl domain, we find that hydrophobic core residues are intolerant to polar substitutions. In the linker, we find residues that are insensitive to swaps to positive charge, hinting at a specific role in ESX function relating to EccB_3_ and EccC_3_. In the transmembrane, we find most substitutions are tolerated apart from specific residues that could play a role in recognition and stabilization of the accessory protein, MycP. Taken together, the observation that most deleterious mutants are in hydrophobic cores or at interfaces implies EccD_3_ is likely primarily a scaffolding protein in the ESX-3 complex.

Our results can also be interpreted in the context of two alternative oligomeric assemblies of ESX systems: dimeric and hexameric^11–14^. Previously, we tagged and purified the ESX-3 complex from the native *M. smegmatis* and observed a dimer as the dominant assembly^11^. We had originally speculated that the EccD_3_ dimer could act as a secretion pore. This is unlikely based on our data, as the transmembrane domain is largely tolerant to mutations in both iron-sufficient and iron-deficient growth conditions. We would expect to see more sensitivity to mutation if the EccD vestibule was translocating substrates, or if interactions with the lipids were required for function. Alternatively, further oligomerization of subcomplexes into a higher-order hexamer with helical rearrangements of core components may enable function. While there are no hexameric structures of ESX-3, there are hexameric structures of the homologous ESX-5 complex. The hexameric structures of the ESX-5 complexes reveal the secretion pore is formed by the transmembrane helices of the EccB and EccC proteins from each subcomplex. The observed LOF effects in our data may suggest the linker hinge motif is sensitive to disruption because of proximity to the central pore formed by EccC and EccB. In the MycP_5_-associated *M. tuberculosis* hexamer, the serine protease MycP_5_ acts as a cap over the periplasmic EccC_5_ and EccB_5_ pore. The EccC_5_ pore is occluded, with the membrane helices rotated to prevent potential translocation of substrates, hinting at a mechanism of secretion where EccD acts primarily as a scaffold to hold proteins in place until some combination of secretion cues enable EccC to rearrange its soluble and transmembrane domains. Additionally, we observed an emergent pattern of effect in the iron-sufficient condition that the MycP bound hexamer structure provides a useful context to explain. The residue with the most dramatic effect was W227, which is on transmembrane helix 4 and extends into a hydrophobic pocket between transmembrane helices 2, 3, and 4. This sensitivity may reflect that hydrophobic residues on transmembrane helix 4 mediate interactions with the MycP transmembrane helix, which has been hypothesized to be required for ESX complex specificity.

We must also consider the caveats of our iron-sensitization assay. We designed our assay to be a negative-selection screen based on the observed phenotype that ESX-3 deletion mutants are less fit relative to wild-type in iron-deficient conditions compared to iron-sufficient conditions. However, we were surprised to see that the variant library behaved similarly when grown in both conditions. This speaks to both the sensitivity of next-generation sequencing and also how that data is scored compared to the individual growth experiments. It is also worth considering that these libraries were screened in a pooled approach, making this screen a co-culture competition, and thus susceptible to trans-complementation. Though it is unclear how ESX-3 function is linked to mycobactin-mediated iron acquisition, it has been shown that ESX-3 mutants are unable to efficiently uptake mycobactin after it is secreted^10,34^. It is likely that ESX-3 secretion specifically is required as the knockout of substrates also results in growth defects in low-iron environments^20,34^. If substrates secreted into the supernatant were responsible for iron-uptake, we expect that this pooled library approach would have revealed functional mutants rescuing the growth of deleterious mutants. It may be that the PE/PPE proteins, which typically remain in the mycobacteria cell envelope, are more directly responsible for cell wall permeability, and thus iron-loaded mycobactin uptake^6,7,34,35^. Furthermore, expression of the ESX-3 operon is regulated by iron such that expression is repressed when there is an internal pool of available iron. As such, we expressed EccD_3_ using the constitutive mOP promoter. Growth in the iron-deficient condition applies selective pressure on variants while also affecting ESX-3 expression, which has been shown to in turn affect function. In our screens, expression of the other complex proteins is regulated by iron availability. This may be especially relevant when we consider the iron-sufficient growth screen, where expression of the ESX-3 complex components is likely reduced in response to the available iron.

In addition to informing about ESX system function, our study is notable for performing a deep mutational scan in *M. smegmatis*, the native organism of the complex. Deep mutational scans are often performed by expressing variants in artificial selection systems in more tractable organisms like *E. coli* or *S. cerevisiae*. Examining proteins in the native context can improve the likelihood of quantitatively measuring mutational effects in the relevant membrane environment and among fleeting interactions. Conducting a pooled mutational scan in *M. smegmatis* is not without difficulty: compared to *E. coli*, mycobacteria have lower transformation efficiency, low frequency of homologous recombination, a genome with high GC content, and grow slower. These constraints limited the synthesis of oligos and successful cloning, resulting in an incomplete EccD_3_ variant library that did not allow us to fully study mutational effects in the transmembrane domain. It would be interesting to study the rest of the vestibule facing residues, as well as the loop between transmembrane helices 6 and 7. In EccD_3-ext_, this loop is near the sensitive linker motif, and also extends into the next protomer.

Our DMS experiments open the door for functional studies of many proteins in *M. smegmatis*, including the other ESX-3 complex proteins to continue studying the secretion cycle. One model is that the available ESX structures represent the complexes in inactive states. We assume activation of the secretion cycle requires the following cues: binding of Esx substrate at the third EccC ATPase domain; the PE/PPE substrates bind and displace the autoinhibitory interaction between the first and second ATPase domain, allowing for ATPase hydrolysis; and multimerization of EccC proteins at the DUF domain to allow for an arginine finger of one EccC to complete the active site of another EccC^36,37^. There may be further rearrangement of transmembrane helices to open the EccC_3_ pore even more, allowing for folded heterodimeric proteins such as the ESX-3 substrates, to travel through^14^. Complete mutational scans of the other ESX complex proteins such as the ATPase EccC_3_ and the serine protease MycP_3_ could reveal hidden patterns related to cues required for oligomerization, secretion pore regulation, and secretion.

Due to the rise of multidrug-resistant and extensively-drug resistant mycobacteria, the development of novel therapeutics is sorely needed. Using DMS approaches in a more relevant model organism would lend itself to mechanistic studies prior to moving to the pathogenic *M. tuberculosis* for drug development. Our mutational scans of the Ubl and linker domains reveal that the larger interfaces between the Ubl domain and soluble domains of other core components are not essential to function making it a poor target for a therapeutic. In contrast, our mutational scan revealed the sensitivity of the linker hinge motif, implying that perhaps targeting the small, electrostatic pocket within the ESX complex could result in an effective therapeutic. More broadly, mutational scanning can identify protein interaction regions essential to growth in mycobacteria that could be targeted with a small therapeutic molecule. Such molecules and approaches could be useful to develop therapeutics against one of the most devastating infectious diseases in the world.

## Methods

### Bacteria strains and media

The *E. coli* strains used for cloning were cultured in LB medium containing 50µg/mL kanamycin. *Mycobacterium smegmatis* strains were cultured in 7H9 medium containing 0.05% Tween-80, and 20µg/mL kanamycin as needed. For the iron-deficient growth experiments, strains were grown in 7H9 medium until late stationary phase, and then washed three times with Chelated Sauton’s medium before diluting in Chelated Sauton’s medium to an OD600 of 0.001. For the iron-sufficient growth experiments, bacteria were grown following the same procedure however they were diluted in chelated Sauton’s medium supplemented with 12.5µM FeCl_3_. Chelated Sauton’s medium contains 60mL glycerol, 0.5g KH2PO4, 2.2g citric acid monohydrate, 4g asparagine, and 0.05% Tween 80 per liter. The pH was adjusted to 7.4 and then the medium was stirred with 10g Chelex 100 resin for 1-2 days at room temperature. The medium was filtered before adding 1g MgSO4⋅7H2O as a sterile solution.

### Cloning

The Δ*fxbA*/Δ*eccD_3_* strain was used for all strain generation. All cloning was performed using the integrative pMV306 vector, which has a both *E. coli* and *M. smegmatis* origins of replication, a kanamycin resistance cassette, and the mycobacterial optimized promoter (mOP). The EccD_3_ complement plasmid was generated via In-Fusion cloning (Takara Bio). The pMV306 vector was linearized via DraI restriction digest. The *eccD_3_* gene was amplified from genomic *M. smegmatis* DNA using oligos designed with 15 bp overhangs that overlapped with linearized pMV306 vector. The point mutants screened for the validation experiment were created via Q5 site-directed mutagenesis kit (NEB) and the pMV306_eccD_3_ vector. Briefly, the NEB base changer was used to design primers such that the forward primer incorporated the new nucleotide sequence in the center of the primer and 10 complementary nucleotides on the 3’ end, and the 5’ end of the reverse primer annealed back-to-back with the forward primer. The template plasmid was removed, and the new variant plasmids were ligated using the KLD mixture provided.

### Library construction

The EccD_3_ variant library was designed to cover amino acids 1-131, which contains the entire ubiquitin-like domain and linker, and residues that were conserved or extended into the EccD_3_ membrane vestibule. Conservation analysis was performed using ConSurf^33^. All structural visualization was performed using PyMol. We designed the oligo library to be inserted into the pMV306 vector, which naturally contains the NotI and NoCI cutsites upstream and downstream the mOP. The oligos contain a NotI restriction cut site followed by mOP sequence at the 3’ end, followed by the entire gene sequence including a single mutation, and a NcoI restriction cut site at the 5’ end. The oligo variant pool was synthesized by Twist Bioscience with codon optimization for *M. smegmatis* expression, and included synonymous mutations and stop codons. The library was received as a lyophilized tube of 5µg DNA, which was resuspended in 100µL 1x TE buffer to a final concentration of 50ng/µL.

We amplified 100ng of insert using 10 cycles of PCR to increase starting material before restriction digests. To prepare insert DNA for insertion into the vector, 1µg of DNA was digested using NotI-HF and NocI-HF for 3 hours at 37°C, and then cleaned using the Zymo DNA Clean and Concentrator-5 kit. Next, 10µg of the pMV306_eccD_3_ vector was digested using NotI-HF, NocI-HF, SacI to remove any residual WT EccD_3_ as there is a cutsite in the middle of the gene, and rSAP to prevent self-ligation of the backbone. The reactions were pooled together and gel extracted (Zymo) to isolate the linearized backbone. The variant oligo pool was ligated into the linearized vector at a 1:3 ratio using T4 ligase overnight at 16°C. The ligation mixture was cleaned and concentrated (Zymo), and eluted using 6µL nuclear free water before electroporation into 50µL MegaX competent cells (Invitrogen). Control transformations using the linearized backbone without insert were performed in parallel. The cells were recovered in 1mL SOC media for 1 hour at 37°C. Then, 10µL recovered cells were used for serial dilutions on LB and kanamycin plates to calculate transformation efficiencies. The remaining recovered cells were added to 50mL LB and kanamycin and grown while shaking at 37°C until the culture reached OD 0.6-0.7. The culture was spun down, and library plasmid DNA was miniprepped (Zymo). Colonies were screened from the serial dilution plates to perform colony PCR and Sanger Sequencing.

### Electroporating the library into *M. smegmatis*

Electrocompetent cells were prepared by growing a starter culture of Δ*fxbA*/Δ*eccD_3_* in 7H9 medium containing 0.05% Tween-80 at 37°C. After 3 days, a 50mL culture of 7H9 and 0.05% Tween-80 was inoculated and grown until OD 0.5. Cells were transferred to a 50mL conical tube and chilled on ice for 30 minutes before harvest at 4000 RPM. Supernatant was removed and cells were resuspended in 25mL ice-cold 10% glycerol. Cells were spun down, resuspended in 10mL ice-cold 10% glycerol, spun down again, and finally resuspended in 5mL ice-cold 10% glycerol. Aliquots of 400µL cells were prepared and used for electroporation immediately, or snap frozen and stored at -80C.

To perform electroporation, 1µg of plasmid DNA was added to 400µL electrocompetent cells. The mixture was added to a chilled 0.2cm electroporation cuvette and placed on ice for 10 minutes. Electroporation was performed using typical electroporation constants for *M. smegmatis*: 2.5kV, 1000W, 25µF. Cells were placed back on ice for 10 minutes, and then recovered in 2mL 7H9 medium at 37°C for 4 hours. 10µL of cells were used to set up serial dilutions for electroporation efficiency calculation. The remaining recovered cells were plated and spread onto a large BioAssay plate (245x245x25mm), and grown at 37°C for 3-5 days. Colonies were scraped when sufficiently large enough to count, however before they were growing onto each other. To scrape, 5mL of 7H9 medium was added to the plate and a scraper was used to collect colonies, and the mixture was collected. Another 5mL of media was added to collect any residual cells. Cells were aliquoted and glycerol stocks were made, snap-frozen, and stored at -80°C.

### Selection experiments

For the variant library selection experiment, growth in both conditions was performed and monitored in parallel, on the same days across all biological replicates. One aliquot of frozen cells was thawed and pelleted, and DNA was isolated to serve as the “time point 0” pre-selection sample (T0). Another aliquot was thawed and allowed to recover in 7H9 medium containing kanamycin until late stationary phase. Cells were then washed three times with Chelated Sauton’s medium before diluting in 30mL Chelated Sauton’s medium to an OD600 of 0.001. For the iron-sufficient growth experiments, bacteria were grown following the same procedure however they were diluted in chelated Sauton’s medium supplemented with 12.5µM FeCl_3_. Three timepoints were harvested every 24 hours across 3 days. 1mL of cells were measured by OD, and then lysed for DNA extraction. After harvest, 1mL of fresh Chelated Sauton’s or Chelated Sautons plus 12.5µM FeCl_3_ was added to the culture.

The point mutant selections were set up and performed in a similar manner, except the time points collected were more frequent. 1mL of cells were collected for OD 600 measurement at 12, 20, 36, 40, and 60 hours. After measurement, 1mL of fresh Chelated Sauton’s or Chelated Sautons plus 12.5µM FeCl_3_ was added to the culture.

### Library DNA preparation and deep sequencing

Cells were lysed by bead-beating with 250μL100μM zirconia/silica beads (BioSpec Products) for 6 minutes. DNA was isolated using a Quick-DNA Zymo Mini-prep kit, and DNA was eluted in 30μL nuclease-free water. Immediately after isolation, all DNA was used for PCR to generate the three amplicons required to sufficiently sequence each EccD_3_ variant. PCR reactions were prepared using the TakaraBio PrimeStar GXL system according to the following recipe: 10μl 5X PrimeStar GXL buffer, 1μl 10μM forward primer (0.3μM final), 1μl 10μM reverse primer (0.3μM final), 10μL DNA, 4μl 10mM dNTPs (2.5mM each NTP), 1μl GXL polymerase, 23μL nuclease free water. The PCR mixtures were amplified with the following thermocycler parameters: initial denaturation at 98°C for 30s, followed by 24x cycles of denaturation at 98°C for 10s, annealing at 60°C for 15s, extension at 68°C for 30s, and a final extension at 68°C for 1 min. PCR reactions were concentrated using Zymo DNA Clean and Concentrator-25 kits. Samples were run on 1% 1x TBE gels to confirm amplicons were successfully amplified.

Libraries were indexed using the IDT for Illumina-TruSeq DNA and RNA UD Indexes. The lengths of the Indexed libraries were quantified using the Agilent TapeStation with HS D5000 screen tape and reagents (Agilent). DNA concentrations were quantified using the Qubit dsDNA HS assay (Invitrogen). All samples were normalized to 4nM and pooled and then paired-end sequenced (V3) on a MiSeq.

### EccD_3_ variant scoring

Demultiplexed paired-end reads were received from the sequencing core as fastq.qz files. The experiment was processed using a snakemake-based pipeline we have developed^38,39^. Each fastq was processed in parallel using the following steps: adapter sequences and contaminants were removed using BBDuk, then paired reads were error corrected with BBMerge and then mapped to the reference sequence using BBMap with 15-mers (all from BBTools)^40^. Variants in the mapped SAM file were called using the AnalyzeSaturationMutagenesis tool in GATK v4^41^. The output of this tool is a csv containing the genotype of each distinct variant as well as the total number of reads. This was then further processed using a python script, which filtered out sequences that were not part of the designed variants, then formatted input files for Enrich2^29^. Enrichment scores were calculated from the collected processed files using weighted least squares and normalized using wild-type sequences. The final scores were then processed and plotted using R. A copy of this processing pipeline, sequencing counts, and fitness scores has been deposited in the Github repositories listed in the data availability section.

### EccD_3_ mutational variant analysis

The EccD_3_ dimer (PDB: 6UMM) was used for structural analysis. We created scores for each type of physicochemistry by defining which amino acids belonged to a given physicochemistry, and then averaging the scores for each missense variant within that group. These scores were mapped onto the B-factors of the structure using the Bio3D R package for visualization.

Conservation of EccD_3_ positions was determined using ConSurf^33^. Specific mycobacterial sequences were obtained from UniProt and the Mycobrowser^42,43^.

Deep mutational scanning data were analyzed in R as described in the text. All scripts used to make figures have been deposited in a Github repository listed in the data availability section.

## Supplemental information

**Figure S1.**
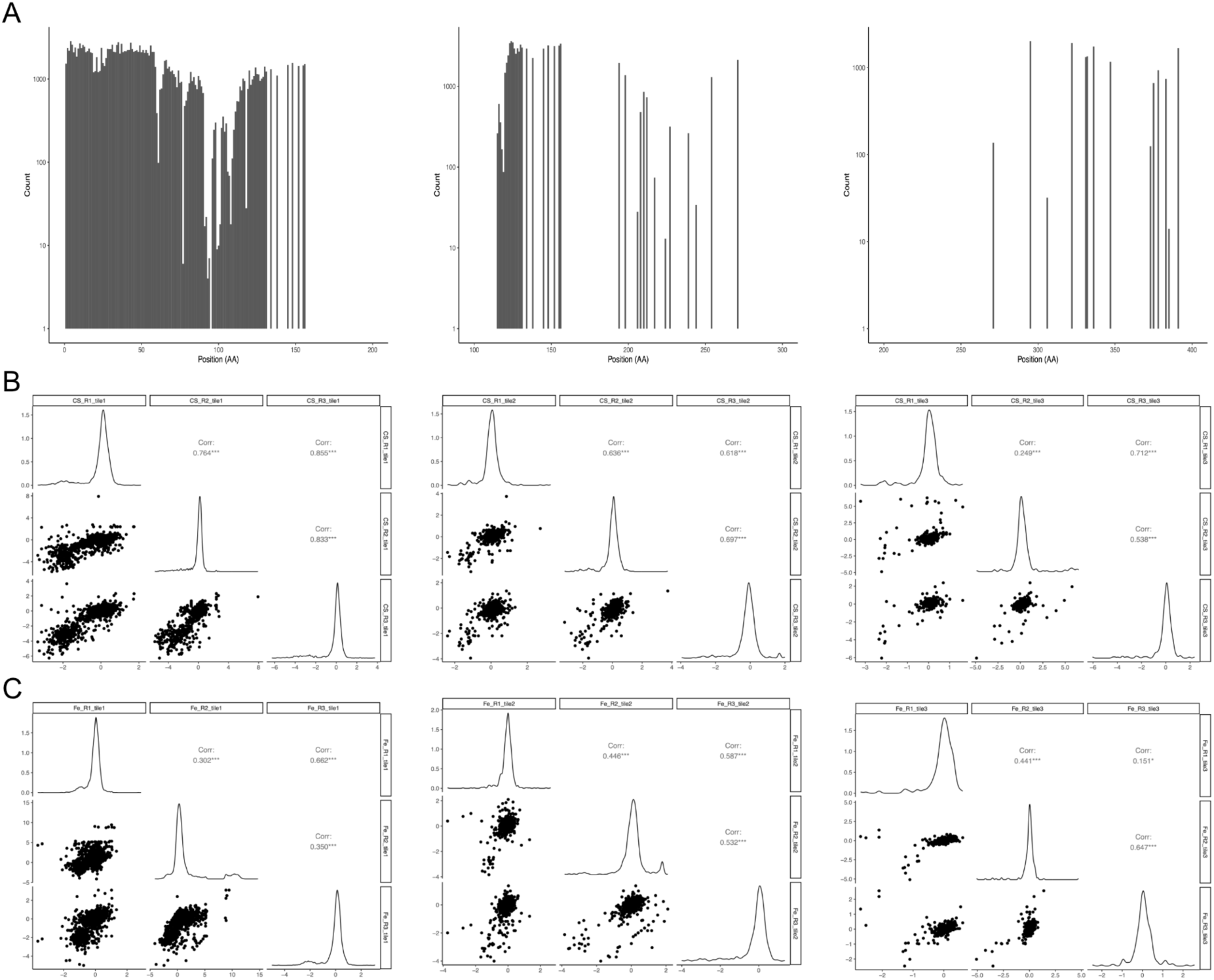
EccD3 library scores and QC. A) Overall distribution of library counts (Y-axis) per position (X-axis) for each tile. B) Cross-correlations between EccD3 iron-deficient screen replicates. Each set of plots represents cross-correlations for each tile. Pearson correlation coefficients above diagonal, histograms of scores for replicates on the diagonal, and dot plots with variants below the diagonal. C) Cross-correlations between EccD3 iron-sufficient screen replicates, same as B.

**Figure S2.**
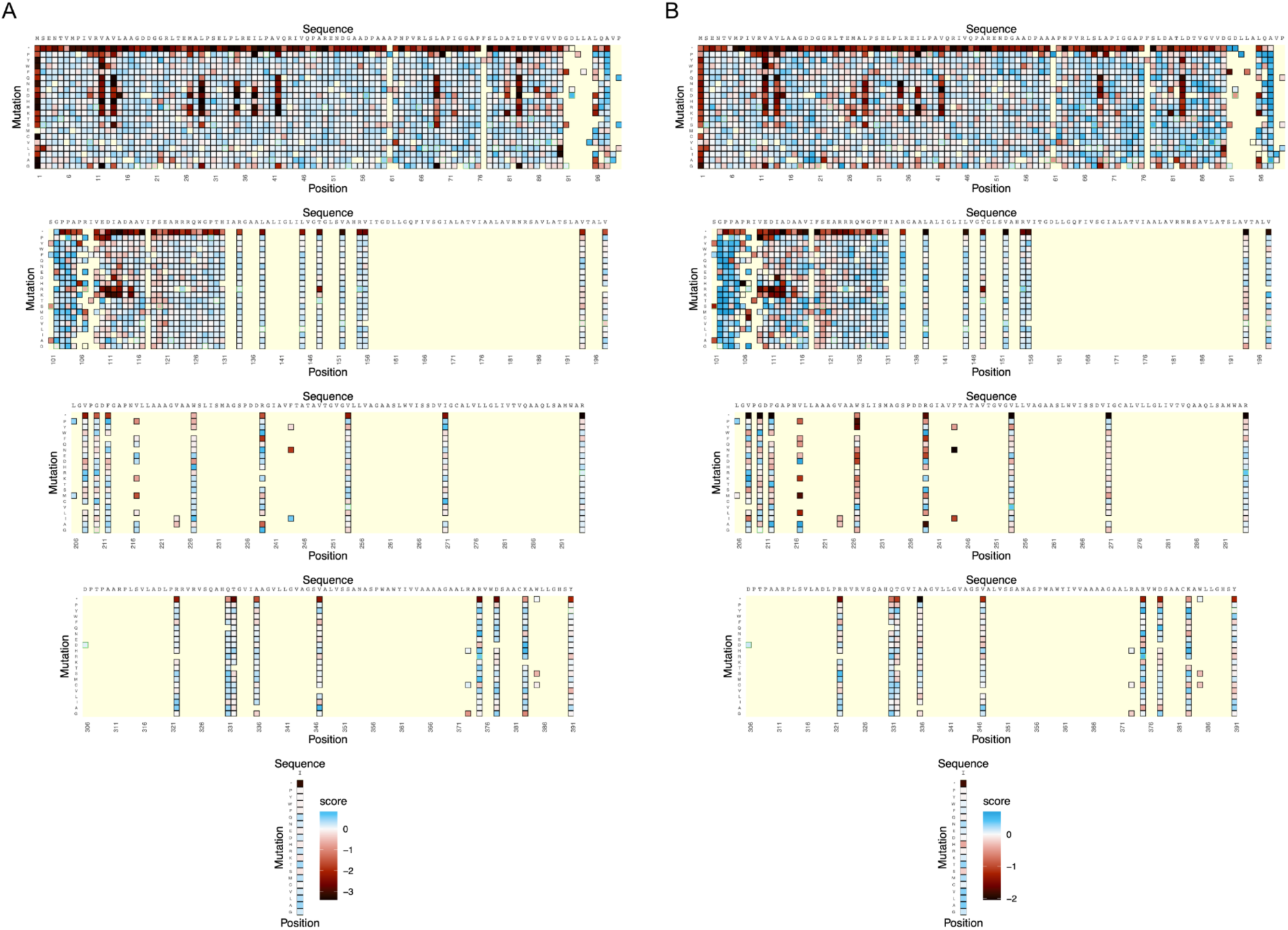
EccD3 heatmaps. A) Heatmap of the EccD3 functional scores from the iron-deficient growth condition. The WT sequence, domain organization, and cartoon secondary structure representation of EccD3 are shown above each section of the heatmap. The variant identity is indicated on the y-axis, and the residue position is indicated on the x-axis. WT-synonymous substitutions are outlined in green and positions not included in the library are light yellow. B) Heatmap of the EccD3 functional scores from the iron-sufficient growth condition.

**Supplemental Table 1.**

| Pre-optimization |  |  |  |
| --- | --- | --- | --- |
|  | Dilution 1, cfu | Dilution 2, cfu | Dilution 3, cfu |
| 100ng | 32 | 3 | 0 |
| 500ng | 67 | 6 | 0 |
| 1µg | 123 | 13 | 3 |
| 2µg | 155 | 49 | 7 |
| Post-optimization |  |  |  |
|  | Dilution 1, cfu | Dilution 2, cfu | Dilution 3, cfu |
| 100ng | 312 | 33 | 1 |
| 500ng | 360 | 63 | 1 |
| 1µg | 647 | 81 | 2 |
| 2µg | -- | 112 | 10 |
| Library 1µg | -- | 285 | 24 |

## Contributions

DDT generated and cloned the EccD_3_ deep mutational scan library. DDT designed and performed deep mutational scan experiments with input and assistance from WCM and OSR. CBM processed raw NGS sequencing data. DDT analyzed deep mutational scanning datasets with input from WCM and JSF. DDT cloned and expressed the site-directed mutagenesis constructs for validation experiments. DDT prepared figures with input from WCM and JSF. DDT wrote the manuscript with edits and approval from all authors. WCM, OSR, and JSF supervised the overall project.

## Conflicts of interest

OSR is an employee and equity holder in GSK plc. JSF is a consultant for, has equity in, and receives research support from Relay Therapeutics.

## Acknowledgements

We thank Eric J. Rubin for providing the Δ*fxbA*/Δ*eccD_3_* and Δ*esx-3* strains. We thank Nadia Herrera for helpful feedback and discussion as we developed and performed these experiments. We also thank Amy Diallo, Babak Javid, the DMS discussion group, and the Coyote-Maestas lab for helpful discussions and feedback as we put the manuscript together. The sequencing was performed by the CZ Biohub SF Genomics Platform. This work was supported by NIH 5F31AI157438 (DDT), NIH 1F32GM152977 (CBM), NIH GM145238 (JSF), and Howard Hughes Medical Institute Hanna Gray Fellowship (WCM), and San Francisco Chan Zuckerberg Biohub (WCM and OSR).

## Data and materials availability

The sequencing data from the EccD_3_ deep mutational scan has been deposited at the NCBI SRA (bioproject PRJNA1151079). Original data files and analysis source code is available at https://github.com/ddtrini/EccD3_DMS.

